# Systematic errors in orthology inference: a bug or a feature for evolutionary analyses?

**DOI:** 10.1101/2020.11.03.366625

**Authors:** Paschalis Natsidis, Paschalia Kapli, Philipp H Schiffer, Maximilian J. Telford

**Affiliations:** Centre for Life’s Origins and Evolution, Department of Genetics, Evolution and Ecology, University College London, London WC1E 6BT, UK; Institut für Zoologie, Universität zu Köln, Zülpicher Straße 47b, 50674 Köln, Germany

## Abstract

The availability of complete sets of genes from many organisms makes it possible to identify genes unique to (or lost from) certain clades. This information is used to reconstruct phylogenetic trees; to identify genes involved in the evolution of clade specific novelties; and for phylostratigraphy - identifying ages of genes in a given species. These investigations rely on accurately predicted orthologs. Here we use simulation to produce sets of orthologs which experience no gains or losses. We show that errors in identifying orthologs increase with higher rates of evolution. We use the predicted sets of orthologs, with errors, to reconstruct phylogenetic trees; to count gains and losses; and for phylostratigraphy. Our simulated data, containing information only from errors in orthology prediction, closely recapitulate findings from empirical data. We suggest published downstream analyses must be informed to a large extent by errors in orthology prediction which mimic expected patterns of gene evolution.

## Introduction

Orthology is a type of homology where the homologous genes originated at a speciation event [1]. The evolution of orthologous genes and the fact that their relationships coincide with species phylogeny, have made them key markers in evolutionary biology. Aligned sequences of orthologs have been used to reconstruct species phylogenies for several decades, but the presence or absence of individual orthologs in genomes of different species is also increasingly being used in various ways to understand evolution - something made possible by the largely complete gene sets now available from genome sequencing projects.

New genes that originated in ancestral species and were passed on to the descendants of this ancestor can be used as synapomorphies of these clades. Matrices recording the presence and absence of sets of orthologs across species have been used to give an estimate of relationships that is assumed to be independent of the traditional sequence alignment-based trees [2]. Gene presence/absence phylogenies of Metazoa have given highly resolved trees showing extraordinary congruence with sequence alignment-based trees [3, 4, 5, 6].

Given a phylogenetic tree, on the other hand, the presence/absence of orthologs in extant taxa can be used to infer gene gain and loss events across their evolutionary history. Such events are being interpreted in the context of origins or loss of key characteristics in those clades. Bursts of gains and losses have been associated with the origins of major animal clades [7, 8] and a search for genes unique to the Bilateria within Metazoa found 157 candidates which the authors linked to bilaterian morphological novelties such as mesoderm and bilateral symmetry [9].

Another use of matrices of gene presence and absence is in phylostratigraphy. Here, genes present in a focal taxon are searched for in the increasingly distant phylogenetic lineages leading to this taxon. In this way it is possible to discover the most distant relatives possessing orthologs and hence to infer the ages of these genes [10, 11]. Sets of genes that may be upregulated in specific developmental stages or structures may have different average phylostratigraphic ages and this information has been interpreted as implying the evolutionary age of traits such as a larval stage [12].

Inferring orthology relationships among thousands of genes that come from distantly related sets of species is a fundamental step in all these studies. Inferring orthologs however, is an inherently difficult task because the genes in a genome evolve in a complex manner [13, 14]. Orthology inference relies on an initial similarity search to identify, amongst all pairs of genes in two organisms, those that are sufficiently similar to be potentially homologous [15]. This step can be difficult if two orthologs have diverged significantly [16, 17].

Subsequent steps are affected by multiple evolutionary processes: genes are frequently duplicated and lost in different lineages; paralogs produced by duplication can evolve at very different rates; and genes can even be transferred horizontally [18, 19, 20]. Our ability to disentangle the relationships between homologous (but not necessarily orthologous) genes is further hampered by the heterogeneities that affect the reconstruction of gene phylogenies such as heterogeneities in evolutionary rates or compositional bias [21].

The three important downstream uses of orthologs we outlined above (presence-absence phylogenies; plotting gene gains/losses across a phylogeny; and phylostratigraphy) must be affected by misidentification of orthologs, but there is little information on the error rates of the methods used to predict orthologs. Importantly, it is not clear what effect inaccuracies in orthology prediction will have on any downstream analyses: are they random with neutral effects or could there be systematic errors in orthology prediction that produce strongly supported results? Part of the difficulty in answering these questions is that, when using empirical data, we do not know the underlying truth.

To overcome this limitation, we have simulated the evolution of sets of orthologous genes along a tree based on the metazoan phylogeny using realistic parameters for sequence evolution derived from empirical data (Fig. 1A). We then used our sets of simulated orthologs to examine the relationship between the frequency of orthology prediction errors and two important aspects of sequence evolution: i) substitution rate and ii) the variance of rate across sites within a gene. Finally, we have explored the effects of these errors on gene presence/absence phylogenies; mapping gene gains and losses on a phylogeny; and phylostratigraphy (Fig. 1B).

**Figure 1.**
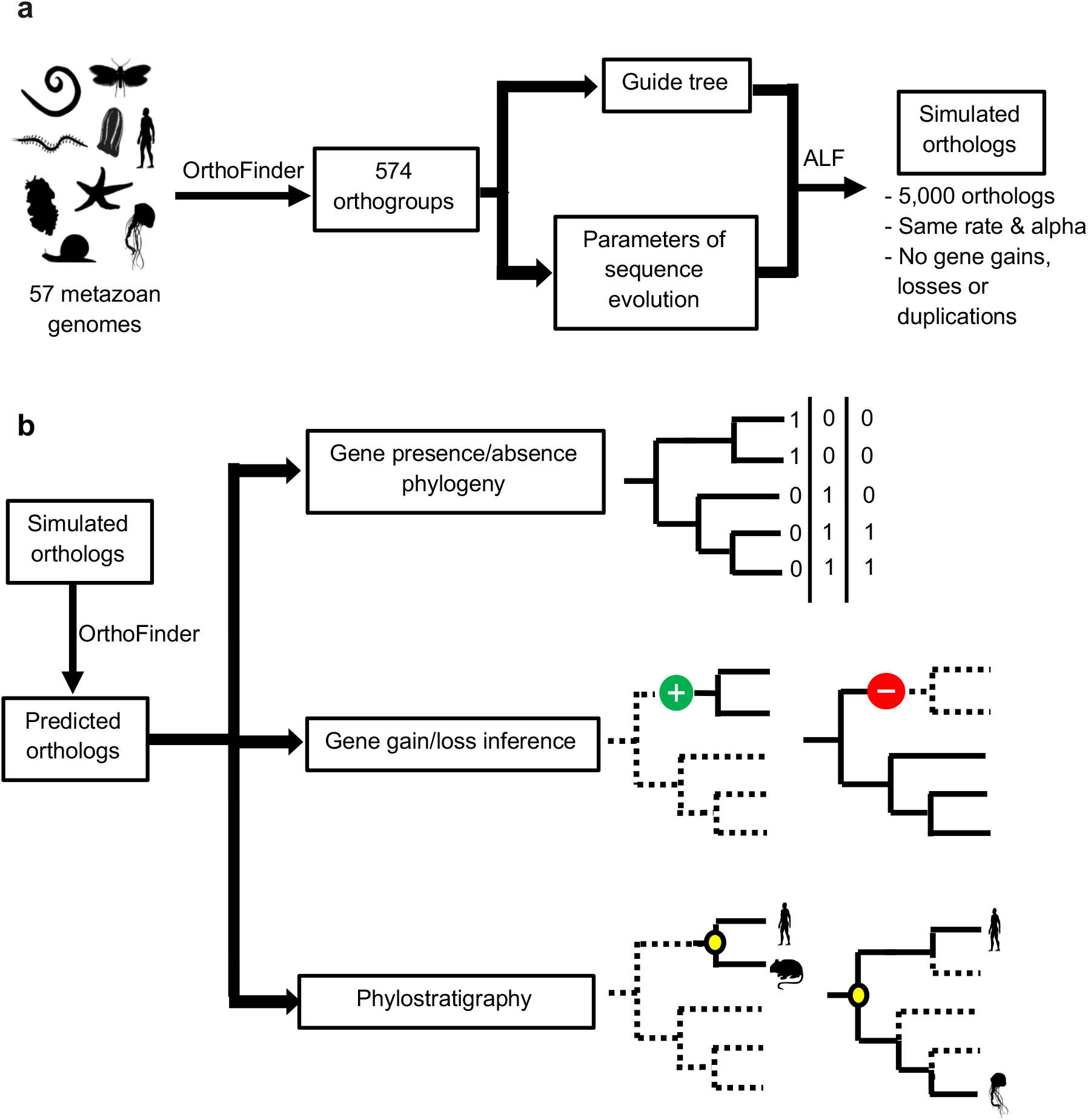
Workflow diagram. **a.** We used information from 574 metazoan orthologs from 57 genomes to infer realistic parameters of sequence evolution to inform our simulations. 200 sets of 5,000 orthologs were simulated according to the empirically derived parameters and a fixed tree topology without any gene gains, losses or duplications. **b.** Orthology relationships among each of the simulated orthologs were inferred using OrthoFinder. These results were used in three different downstream analyses to understand the impact of orthology prediction error: gene presence/absence phylogeny; gene gain/loss inference; and phylostratigraphy.

## Results

### Estimating simulation parameters from empirical data

We chose to simulate the evolution of sets of orthologous genes across a tree based on the relationships among metazoan phyla - a part of the tree of life that has been repeatedly studied in the ways we describe. Basing our simulations on this real example allows us to interpret how different rates of evolution across lineages might affect orthology inference, enables us to use realistic parameters for the simulation of evolving genes and means we can compare results and conclusions from real data with the results of our simulations. In order to isolate the effects of errors in orthology prediction, we simulated with no gains or losses of genes and no duplications. For analyses using the presence or absence of orthologs predicted from our simulated data, the only signal present comes from errors in orthology prediction.

The guide tree (Fig. 2) on which we based our simulations contained 57 species from across the Metazoa. We fixed its topology based on the current consensus for the relationships between animal phyla [22–25]. We built a concatenated alignment of 574 near-global groups of metazoan orthologous genes that we had predicted from our 57 genomes using OrthoFinder [26] with default settings. The branch lengths of the guide tree were calculated using IQ-tree [27] using the LG+F+G+C60 model and our concatenated alignment.

**Figure 2.**
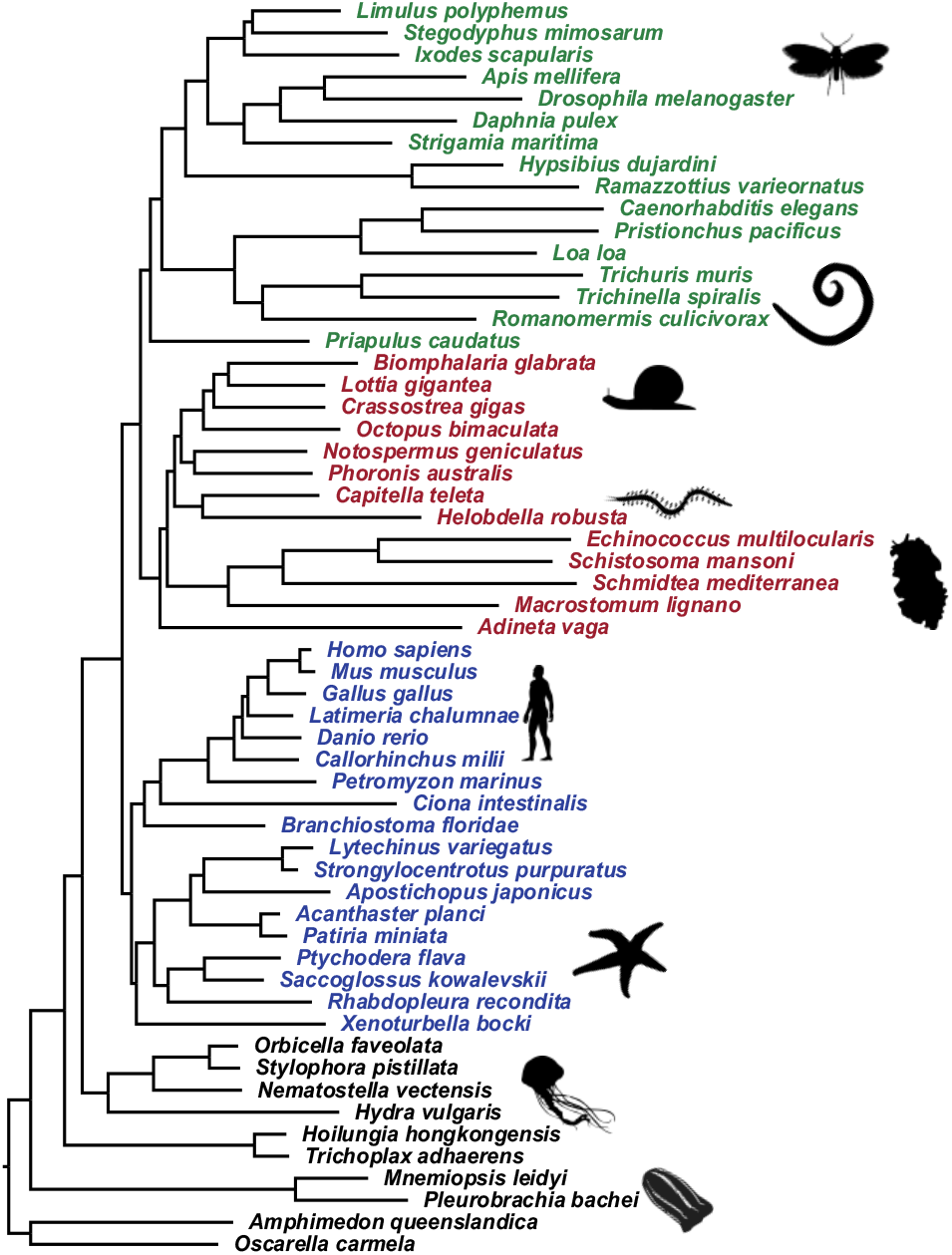
Guide tree used for all simulations. The guide tree under which the orthologs evolved in our simulations. Branch lengths were estimated based on the concatenated set of 574 orthogroups using the LG+F+G+C60 model. Each simulation involved the evolution of 5,000 orthologs along a scaled version of this guide tree, where all branch lengths were multiplied by a scalar ranging from 0.2x to 10x.

In the context of our fixed topology, we used each of the 574 genes individually to estimate 574 gene-specific sets of parameters of the evolutionary model: amino-acid frequencies; branch lengths; and the alpha parameter for rate variation among sites. These measurements were then used to build distributions of these parameters (normal distributions for alphas and branch lengths, Dirichlet for amino-acid frequency vectors).

### Simulating sets of orthologs

We randomly sampled values from the distributions of parameters and applied these to make our simulations as realistic as possible. We performed 200 simulations using the ALF software [28]. Each simulation involved the evolution of 5,000 orthologs along the guide tree following the amino-acid exchangeability values of the LG model.

As the 574 genes we chose were selected on the basis of being found in the majority of taxa, they are likely to be a biased sample that does not represent the real evolutionary diversity of genes. To account for this and to obtain a more realistic diversity in evolutionary rates in particular, in each simulation, we multiplied the guide tree length by a gene rate multiplier (the same one for all 5000 genes within a simulation). With a higher gene rate multiplier the sequences accumulate more mutations through the simulation thus emulating faster-evolving genes; with a lower gene rate multiplier, the simulated genes evolve more slowly. The gene rate multipliers ranged from 0.2x (5 times slower) to 10x (10 times faster). The larger multipliers effectively extended the rates considered well beyond those represented by the 574 genes selected during our conservative ortholog selection. For each simulation, we also selected at random a single alpha parameter for rate variation among sites from the normal distribution of alpha values. This alpha parameter was applied to all 5000 genes simulated for that simulation.

### Effects of gene rate and between-site rate heterogeneity on OrthoFinder accuracy

For each of the 200 sets of simulated proteins (for which we know the gene rate multiplier and alpha) we ran Orthofinder using default settings. With perfect orthology prediction, we would expect to recover exactly 5,000 orthogroups and each orthogroup would contain exactly 57 genes, one for every species. Any divergence from these numbers will be due to orthology inference errors.

In Fig. 3 we show the relationship between both the gene rate multiplier and the alpha parameter and the numbers of predicted orthogroups and mean orthogroup sizes. In the simulations with small gene rate multipliers, representing slowly-evolving genes, OrthoFinder was very successful in recovering the correct number of orthogroups. As the gene rate multiplier increases and the underlying genes become faster evolving, however, we see that OrthoFinder makes increasing numbers of errors and that the predicted number of orthologs becomes higher than 5,000. In simulations with the highest gene rate multipliers we see as many as 250,000 orthogroups and mean orthogroup size reaching values around 2.5 genes/orthogroup; is much smaller than the true size of 57.

**Figure 3.**
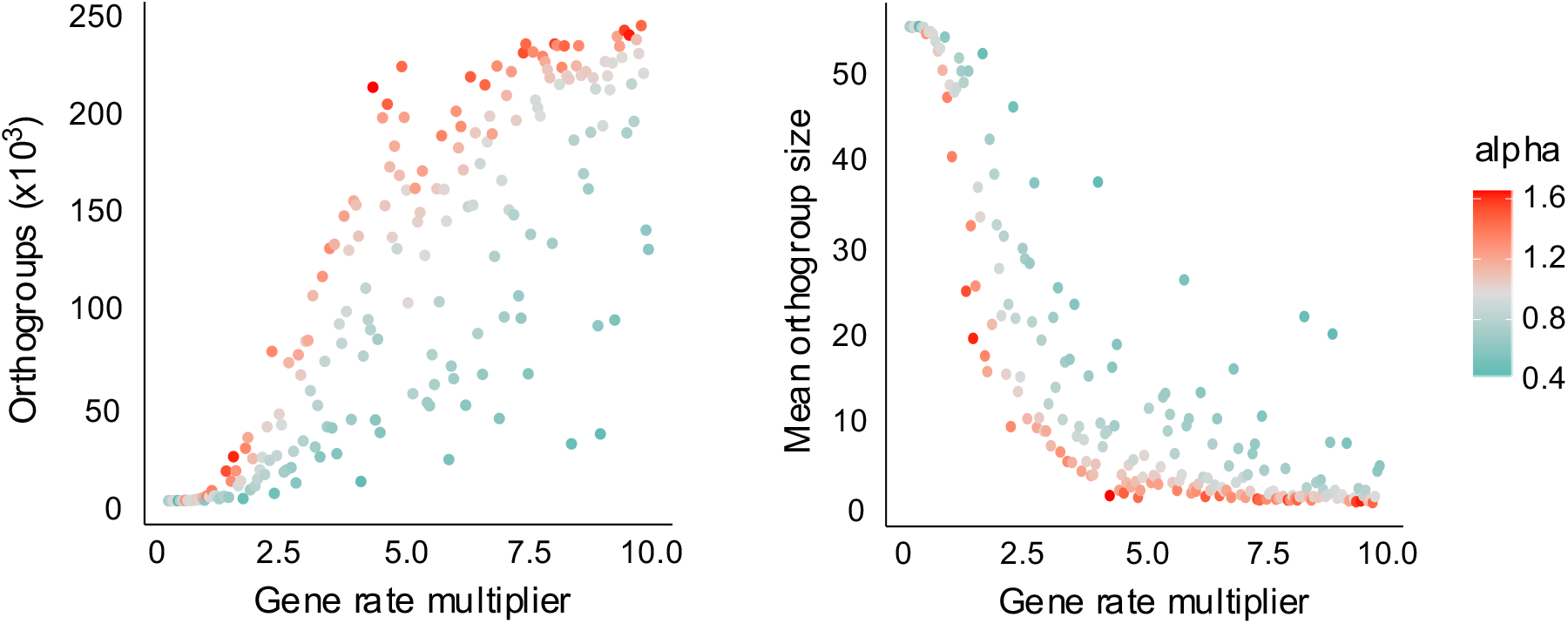
Errors in orthology prediction are more frequent with faster genes and with higher alphas. Number of orthogroups inferred from each of the 200 simulation replicates plotted according to rate of evolution and alpha. An accurate inference would contain 5000 orthogroups (left). Mean orthogroup size inferred from each of the 200 simulation replicates plotted according to rate of evolution and with different alphas (right). An accurate inference would show orthogroups containing 57 species. Higher orthogroups sizes indicate more errors. In simulations with small gene rate multipliers (corresponding to slow-evolving genes) orthology inference was successful in recovering 5,000 orthologs with the correct mean size of 57 genes. With larger gene rate multipliers, orthology inference erroneously inferred more and smaller orthogroups. Higher alphas (less between-site rate heterogeneity) result in more errors in orthology inference.

We wanted to see whether the relationship between rate of evolution within an orthogroup and the number of species included that orthogroup is also reflected in empirical data. To control for phylogenetic distance we considered the sets of orthologous genes in common between 4 separate pairs of species. For each pair of orthologous genes we measured the patristic distance between the pair of species and compared this with the total number of species represented in that orthogroup. We found across all four comparisons that, just as with our simulated data, there was an inverse correlation between rate of evolution of a gene and the number of species in the orthogroup containing that gene (Supp. Fig. 1).

We also see that the alpha parameter for rate variation among sites has an independent effect on the size of error. For a given gene rate multiplier, higher values of alpha (less between-site rate heterogeneity) leads to more errors. While gene rate may be the principal driver of errors in orthology inference, the degree of site rate heterogeneity also seems to be an important factor.

The higher frequency of errors with increasing gene rates is not unexpected. As orthologs become more divergent, it becomes more difficult to determine their homology. To use *reductio ad absurdum* - two genes at either end of an infinitely long branch will have sequences as different from each other as random sequences. The effect of the higher alpha parameter on our ability to infer orthologs correctly is less easily explained, however, since this affects only the distribution of rates across sites (higher alpha parameters have more uniform rates) but not the mean rate. The low alphas (more skewed rates across sites) mean that there are many slowly evolving sites and a few fast evolving sites. We speculate that the presence of sufficient numbers of slow evolving sites would permit the similarity searching stage of orthology prediction to find regions that are similar enough for the genes to be considered as homologs. For genes with higher alphas (a more even distribution of intermediate rates) if the rate across the whole gene is high enough, then the similarity search may fail to find any regions with sufficient similarity to warrant further consideration by the Diamond BLAST algorithm [29].

### Gene presence/absence phylogenies are informed by errors in orthology inference

The increased availability of well-characterised metazoan genomes has allowed the presence or absence of orthologous genes in different species to be used as characters to reconstruct phylogenetic relationships (e.g. metazoan phyla [3–6] and Insecta [30, 31]). The assumption underlying these studies is that the 1s and 0s of the matrix represent real presences and absences of genes within genomes and hence gains and losses of genes through evolution. These presence/absence phylogenies are highly congruent with sequence based metazoan phylogenies [22–25]. It is not clear, however, whether the phylogenetic signal contained in the gene presence/absence matrices might be affected by orthology inference errors and how these errors might influence the resulting tree.

With our simulated data we have observed that there are frequent errors in orthology prediction which increase with faster evolving genes (Fig. 3) and with higher alphas. We wanted to know what effect these errors might have on our ability to reconstruct gene presence/absence phylogenies. The inferred sets of orthologs from each of our simulation experiments constitute a matrix of presence and absence of orthologs immediately suitable for phylogeny reconstruction. Since we simulated without any gene gain, loss or duplication, any phylogenetic information in the matrix will come solely from orthology prediction errors.

For each simulation replicate we used the corresponding gene presence/absence matrix to build a phylogenetic tree using RAxML [32] with the BINGAMMA model as appropriate for the evolution of binary characters. To evaluate the ability of each simulated matrix to inform tree reconstruction, we used the Robinson-Foulds (RF) distance to measure the difference between the true tree relating the taxa (the simulation guide tree) and each tree built using the orthologs inferred from our simulated data (Fig. 4A). RF measures the number of splits that differ between the tree inferred using simulated data and the guide tree. RF is 0 if trees are identical and 1 if they are completely different.

**Figure 4.**
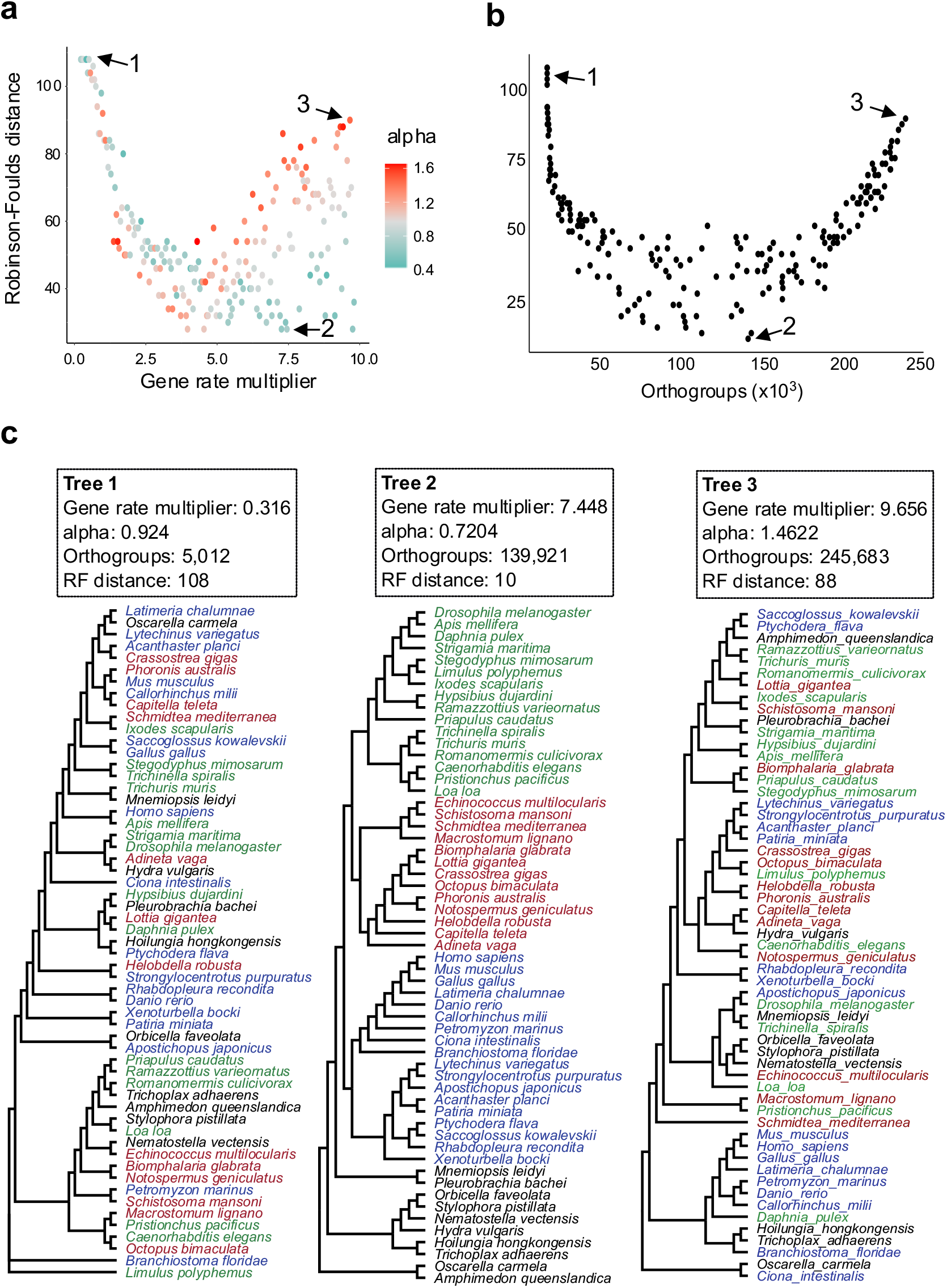
Gene presence/absence phylogenies benefit from errors in orthology inference. **a.** Relationship between gene evolution rate on the accuracy of trees reconstructed from the per-species presence//absence matrix for each simulation. Accuracy is calculated using the Robinson-Foulds distance (RF) between the true tree and the reconstructed tree. In simulations of slow-evolving genes (few orthology inference errors) the corresponding presence/absence trees are very poor (High RF). For faster-evolving simulations (more orthology inference errors) the trees become much more accurate. For slower genes a higher alpha gives better trees. As the rate increases a lower alpha results in superior trees. The values corresponding to the trees (1,2,3) shown in part C are indicated by arrows. **b.** The most accurate trees correspond to an intermediate level of error as measured by the number of inferred orthogroups. With very low and very high error rates the trees are very poor. The values corresponding to the trees (1,2,3) shown in part C are indicated by arrows. **c.** Examples of trees reconstructed using matrices of gene presence/absence based on slow, intermediate and fast evolving simulations. The trees correspond to the points indicated by arrows in Figure parts A and B. The parameters used in the three simulations are indicated in the boxes. Green species are ecdysozoans, brown species are lophotrochozoans, blue species are deuterostomes.

Not surprisingly, slowly evolving genes, where most or all orthologs were correctly predicted, contained little or no phylogenetic information. The trees estimated using these data were essentially unresolved giving a high RF. As we considered faster-evolving sets of genes, we saw the appearance of phylogenetic structure in the trees as shown by a large decrease in RF values. The best trees (lowest RF) were observed in simulations with higher rates but when alphas are low. Higher rates with higher alphas perform poorly. The best trees (lowest RF) actually correspond to an intermediate rate of orthology inference error as can be seen in Fig. 4B. Examples of trees from the extremes of this distribution (very few or very many errors) illustrate the poor estimates of phylogeny when compared with the best trees we observe at intermediate levels of error (Fig. 4C). Importantly, the orthology errors caused by high substitution rates and small alphas are far from random, they contain information that accurately reflects the underlying species relationships.

To compare our best trees from simulated data with one built using real presence/absence data, we built a gene presence/absence matrix using orthogroups predicted using OrthoFinder on our sets of genes from the same 57 species. We reconstructed a phylogeny from these data using MrBayes [33] with ascertainment bias correction for singletons and all-absence sites (Fig. 5A). We compared this real presence/absence tree with the best-scoring presence/absence tree from our simulations (gene rate multiplier = 7.448, alpha = 0.7204). We found that the tree generated using real data (that includes real gene presence/absence signal) and the tree from our simulations (resulting purely from orthology errors) are highly congruent. The results suggest that much of the phylogenetic signal in the real gene presence/absence matrix may be derived from errors in orthology inference rather than from real gene gains and losses.

**Figure 5.**
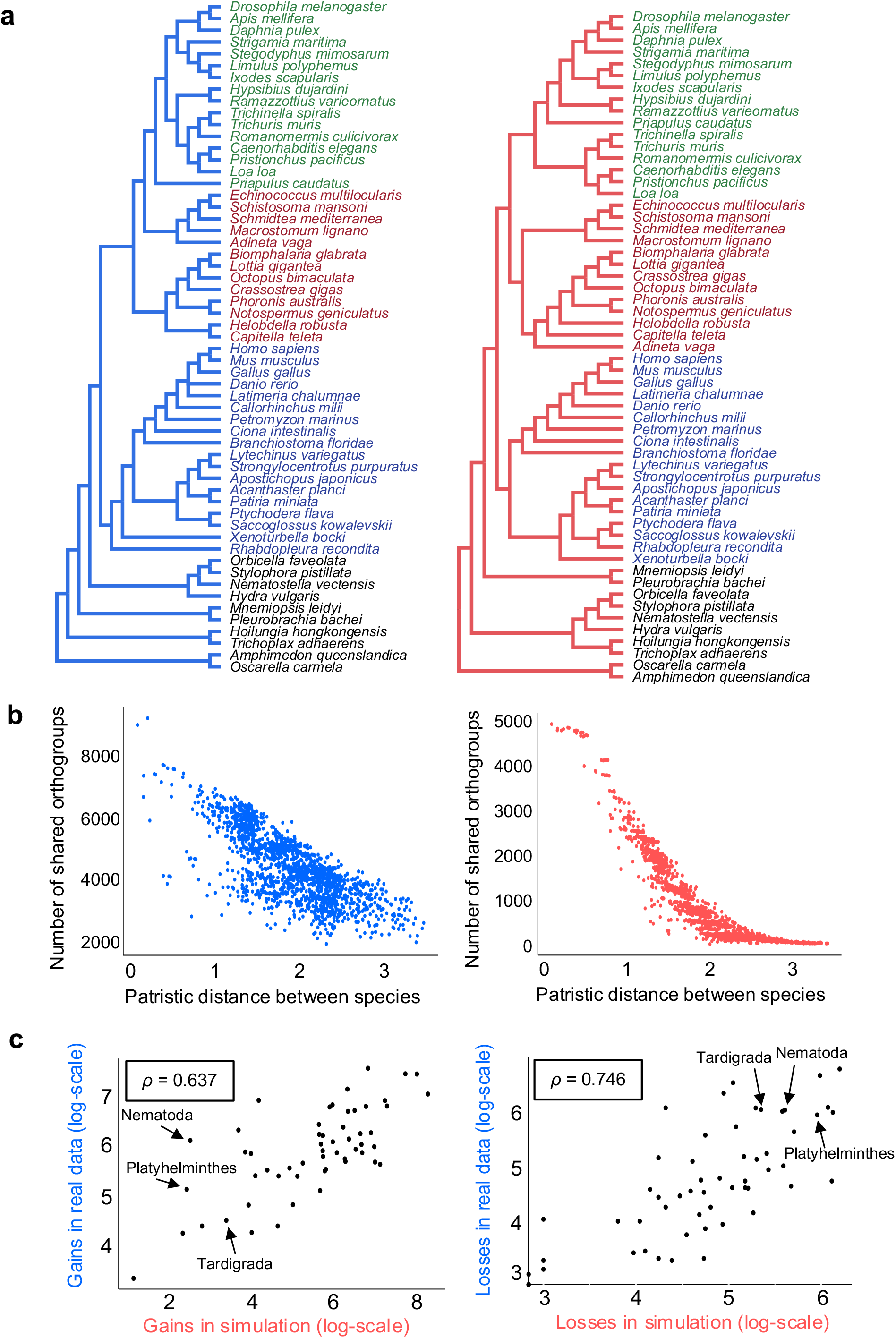
Downstream analyses based on orthology prediction errors in simulated data closely resemble the results from real data. **a.** A phylogenetic tree reconstructed using the gene presence/absence matrix from real data (left/blue) closely resembles the tree based on orthology prediction errors from simulated data (right/red). Green species are ecdysozoans, brown species are lophotrochozoans, blue species are deuterostomes. **b.** The number of orthogroups shared between pairs of species correlates with the patristic distance between them for orthology predictions based on both real data (left/blue) and for simulated data for which all information results from errors (right/red). **c.** Comparison of numbers of gene gains and losses in each node of the guide tree estimated from real (y axis/blue) and simulated (x axis/red) data. Numbers of gene gains in each node (left) and gene losses (right) are strongly correlated between simulated and real data. Each dot represents an internal node of the guide tree. The values for the nodes leading to the fast evolving tardigrades, platyhelminths and nematodes are indicated. The correlation coefficient ρ was calculated using Spearman’s rank test.

We have found that, at least for data sets with intermediate levels of orthology inference errors, the set of orthologs that any given pair of species have in common reflects their phylogenetic relationships; we infer that the number of orthologs in common to any given pair of species would therefore be related to their evolutionary distance. To test this, for all possible pairs of species in the guide tree (Fig. 2), we plotted the patristic distance between them (the sum of branch lengths separating them) against the number of inferred orthogroups they share. We did this both for the real data and for the simulation that had resulted in the most accurate tree (gene rate multiplier = 7.448; alpha = 0.7204). For both real and simulated data (Fig. 5B) we see that there is a strong negative correlation between the number of common orthologs and the evolutionary distance between species.

Our results show that the errors in orthology are far from random, but are strongly correlated with phylogenetic relationships of the species in question and the degree of similarity among their orthologs. Faster-evolving genes are in general less likely to be correctly grouped as orthologs. We find that genes in pairs of species that are evolutionarily distant are also less likely to be correctly identified as orthologs. These errors are exaggerated when the alpha parameter for site-specific rate variation is sufficiently large.

### Numbers of gene gains and losses are systematically overestimated due to orthology inference errors

The ability to work with complete sets of genes within genomes has prompted efforts to infer the series of gene gains and losses that occurred along the internal branches of the evolutionary tree relating a given set of species [7, 8]. This approach is seen as a way to uncover possible genomic correlates of important phenotypic transitions in evolutionary history: the evolution of clade-specific novelties; losses of certain morphological characteristics; or appearance of certain embryological characters [12]. One inference from recent work reconstructing gene gains and losses across the animals has been that the evolution of the metazoan gene repertoire has been driven to a great extent by gene loss events [7], with losses especially prominent in the branches leading to some fast-evolving animal phyla (Nematoda, Tardigrada, Platyhelminthes) [8].

We have repeated a similar analysis using a parsimony approach to map gains and losses of genes onto our guide tree [34]. We counted the numbers of gains and losses at different nodes of the tree reconstructed using our presence/absence matrix derived from real data. We compared these results with those from one of our matrices derived from simulated data.

For each internal branch of the guide tree, we compared the number of gene gains and losses inferred using real data and from the matrix of orthologs derived from the simulation that gave the best-scoring presence/absence tree (see Fig. 4). We observe a strong correlation between the numbers of gains and losses reconstructed at each node using real data and data simulated with no gene gains or losses (correlations: gains ρ=0.637; losses ρ=0.746. Fig. 5C). The similarity between real data and our simulations suggests that some of the apparent gene gains and losses in analyses of real data are likely to be due to systematic errors during orthology inference related to the distance between taxa.

### Apparent gene ages in phylostratigraphy analyses are correlated with rates and phylogenetic distance

Phylostratigraphic analyses estimate the ages of each of the set of genes in a focal species by looking for their homologs in increasingly distantly related sister clades. The most distant outgroup species or clade in which a homolog is found defines the age of the gene. Simulations have been used previously to assess the accuracy of phylostratigraphy and have shown an inverse correlation between rate of evolution and apparent gene age [35, 36, 37].

For each of our simulated sets of orthologs, we considered *Homo sapiens* as the focal species and, for each orthogroup containing a human gene, we found the most distant sister clade also contained in the orthogroup to assign the human gene to a phylostratum. In our sets of simulated proteomes, all genes originated at the root of the tree. Each human gene was assigned a value corresponding to the taxon present in the corresponding orthogroup most distantly related taxon to human (Fig. 6A). For each simulation we calculated the average value of the 5,000 human genes to give an Average Gene Age (AGA). We plotted the AGA in relation to the gene rate multiplier and the alpha of each simulation (Fig. 6B). Under perfect orthology prediction we would expect all human genes to be classified as metazoan genes with AGA equal to 1. For simulations of slowly evolving genes the AGA is, as expected, close to 1 (most/all genes originating in the metazoan ancestor). For simulations with faster evolving genes, the AGA steadily increases meaning that the 5000 human genes appear to be increasingly young.

**Figure 6.**
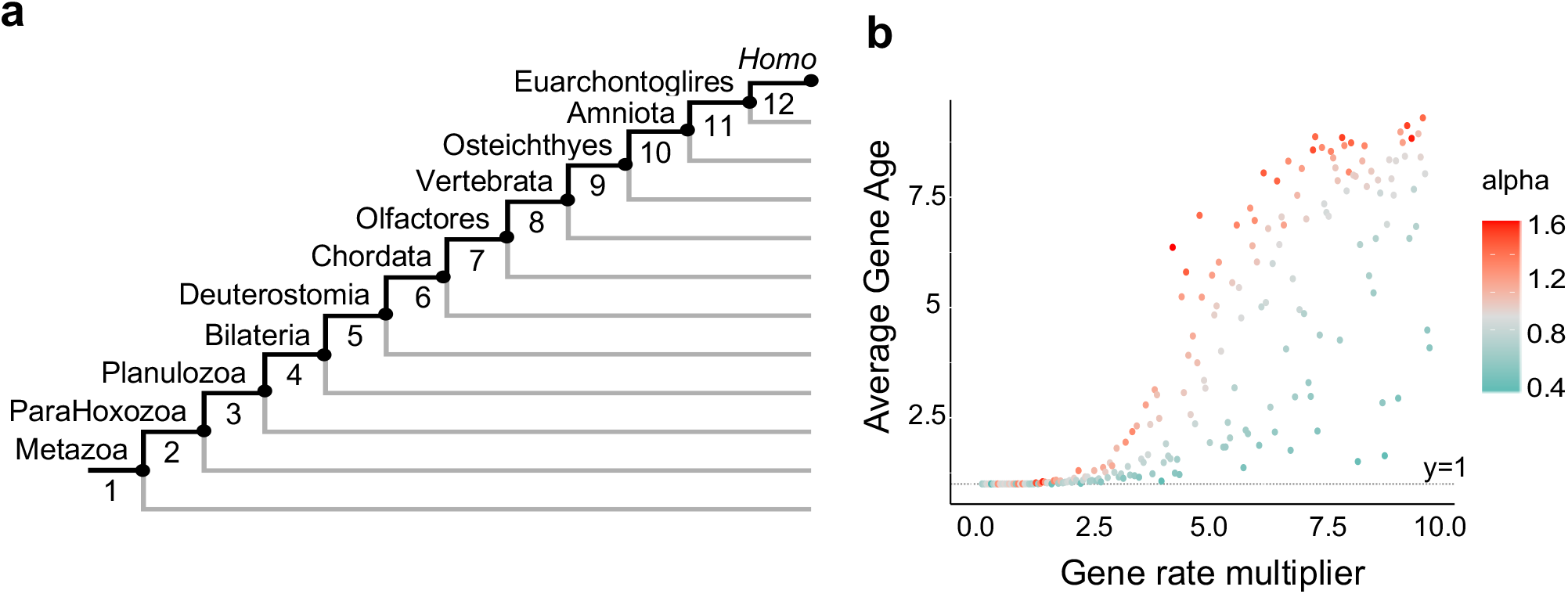
Fast evolving genes appear younger than they are in phylostratigraphic analysis. **a.** The phylostrata used to calculate the age of each human gene in each simulation replicate. The numbers at each node were used to calculate the Average Gene Age (AGA). **b.** Average gene age scores for the 200 simulation replicates. The score represents the average phylostratum value for each of the 5,000 human genes in the simulation. The expected score with no orthology inference errors would be 1.

## Discussion

Sets of predicted orthologs are being used in different ways for several important analyses of evolution. Evolutionary trees based on the presence/absence of orthologs (and homologs) are strikingly similar to those derived from sequence alignments and have been taken as important corroborative evidence of phylogenetic relationships. Mapping the distribution of predicted orthologs onto a known tree is being used to draw conclusions about the genomic correlates of the emergence of major clades and the gene gains and losses associated with these important evolutionary events. Phylostratigraphic analyses of gene ages, likewise, are founded on the assumption that orthologs in different species have been accurately identified.

Correct orthology identification is difficult [13, 14], especially between more distantly related species - genes inevitably become more distinct as they diverge [16]. But an implicit assumption of the downstream analyses of sets of orthologs is that, while errors in identifying orthologs are to be expected, there should be no systematic biases in the distribution of the errors that will be interpreted as signal in subsequent steps.

We have shown that errors are not randomly distributed when we consider realistic simulated sets of orthologs and we suggest that this problem will also affect real data. While the likely artificial enhancement of the phylogenetic signal might be seen as a benefit in the case of presence/absence phylogenies, conclusions drawn concerning the gains and losses of genes on a phylogeny may well have been based on a misleading signal derived from the systematic biases we have identified. Phylostratigraphic analyses will be similarly affected, with fast evolving genes appearing to be younger than their real age, as has been pointed out previously [35, 37].

It is not immediately obvious how to separate the signal derived from gains and losses that are to be expected of a real evolutionary process from apparent gains and losses due to errors in orthology inference. The apparent gains and losses that our simulations predict will follow a very similar pattern of distribution expected of real events. This problem is compounded by the fact that we cannot know the true pattern of gains and losses. Our simulations, although missing important aspects of the process of gene evolution, could be used to derive a rough null expectation of the degree of error that might be subtracted from the total signal. Branches with numbers of gains and losses strikingly in excess of the null expectation are likely to represent a real signal indicating a spike in gene gains or losses, for example. Ultimately, however, we need to aim to reduce the errors as far as possible, meaning that extra steps to correct for errors correlated with evolutionary distance are required. The large effect of different distributions of between-site rate heterogeneity (alpha) on the frequency of error could also be an interesting new avenue of research that may help refine orthology prediction.

We have shown that each of the results we derived from simulated data can mirror observations from empirical data. While some of the evolutionary signal found in the sets of orthologs derived from empirical data must represent real events of gene gain and loss, our results suggest that this signal is likely to be supplemented to an unknown degree by the systematic errors we have described.

## Methods

### Extracting simulation parameters from empirical data

We collected the set of predicted proteins from 57 well-characterised metazoan genomes selected to give a broad representation of the different metazoan clades (10 non-Bilateria, 9 Xenambulacraria, 9 Chordata, 13 Lophotrochozoa, 16 Ecdysozoa) and exhibiting different rates of evolution.

We used a consensus of the current status of metazoan phylogeny [22–25] as a guide tree for our simulations of gene evolution. To get estimates of branch lengths from real data, we inferred orthology relationships among the 57 proteomes and selected 574 orthogroups that contained orthologs from at least 46 (i.e. >80%) species and with a maximum of 4 gene copies per species. The selection was performed using the custom script OGFilter.

We aligned the 574 sets of orthologs with MAFFT [38], trimmed them with BMGE [39] (settings: -h 0.7 -g 0.7) and used these individual gene alignments to build trees with iqtree v1.6.12 (LG+G+F model). Likely paralogs were then deleted from the set of orthologs based on these gene trees using a custom script ‘ParaFilter’. The resulting 574 single-copy orthogroups were then realigned, concatenated into a single matrix of 210,516 aligned amino acids and this was used to run an MCMC chain in PhyloBayes with the CAT-Poisson model.

We used the 574 single-copy orthogroups and the fixed guide tree (iqtree -g command) to estimate the following parameters of the LG model of sequence evolution: amino-acid frequencies, gene tree lengths (sum of branch lengths) and alpha parameters for rate variation among sites. These sets of parameters were subsequently fitted into distributions (Dirichlet for amino acid frequencies, normal for alphas), that were later used to provide realistic parameter values for the simulations. The tree lengths were used to define guide tree multipliers (gene rate multipliers) in order to simulate slower and faster evolving genes. The multipliers obtained from the 574 orthogroups ranged from 0.2x to 3x, but we extended the range up to 10x in order to capture genes with evolutionary rates that were presumably missed during our stringent approach for selecting orthogroups.

### Orthology inference

We inferred orthology relationships among the simulated sets of orthologs using OrthoFinder v2.3.1 [26] using the default settings. We did one orthology inference per simulation repeat. From the OrthoFinder output, we counted the number of resulting orthogroups and the mean orthogroup size. Since each simulation repeat was run with a specific guide tree length and a specific alpha parameter for rate variation among sites, we were able to correlate these two parameters with the number of orthology errors that we observed.

### Running simulations of sets orthologous genes

We performed our simulation experiments using ALF [28] under the guide tree and using parameter values derived from real data as described above that were provided in a replicate-specific configuration file. We created 200 ALF configuration files with settings for the guide tree lengths (guide tree in Fig. 2 with all branch lengths multiplied by a scalar between 0.2x and 10x), amino-acid frequencies and alpha parameter for rate variation among sites were chosen at random from the empirically derived distributions described above. Each configuration file produced one simulation repeat. Each repeat was run with 5,000 starting genes, 100 possible amino-acid frequency states and a single alpha value of the gamma distribution to model rate variation among sites. Each of the 5,000 genes was evolved along the guide tree according to the LG matrix exchangeabilities and independent from other genes. We did not allow for any gene losses or duplications to occur during the simulated gene evolution. As a result, at the end of each simulation we had 5,000 sets of orthologs present in a single copy in all 57 species.

### Gene presence/absence phylogeny inference

We converted the Orthogroups.GeneCounts.csv matrix from the OrthoFinder result to a gene presence/absence binary alignment using a custom script (orthocounts2bin). This script creates a FASTA or PHYLIP alignment from the gene count matrix where every non-zero character is coded as 1 and every zero character is coded as 0.

Since the gene count matrix does not contain information for the unassigned orthologs (singletons, ‘orphan’ genes), we added these to the gene presence/absence binary alignment using a custom script. The resulting per-species ortholog presence/absence matrix (singletons included) was used to infer gene presence absence phylogenies using RAxML [32] using the ‘-m BINGAMMA’ model. No ascertainment bias correction was used since singletons are present in the alignment. We also reconstructed a tree using the gene presence/absence information in the real sets of genes from the 57 species. We used MrBayes [33] with the F81-like model for binary data and using the ascertainment bias corrections ‘nosingletonpresence’ and ‘noabsencesites’.

#### Gene gain and loss inference

We used a parsimony optimisation approach in PAUP* [33] to infer gene gain and loss events in each internal node of the guide tree. We converted the 200 gene presence/absence matrices into nexus scripts suitable for PAUP* input using a custom python script. We then ran the data matrix with the script in PAUP* for each of the 200 simulations and parsed the PAUP* output to infer the number of gene gain and loss events that occurred on each internal node. We did the same for a per-species ortholog presence/absence matrix derived from the real sets of genes from the 57 species.

#### Phylostratigraphy analysis

We used the OrthoFinder results from our 200 simulated sets of proteomes to examine the effect of orthology error on phylostratigraphic analyses of gene age. We chose the human as focal species, and looked at the 5,000 orthogroups that contained a human gene. We assigned an age to each of these orthogroups based on the species it contained, according to Fig. 6A. Each simulation received an average gene age score that is the average age over all orthogroups that contained a human gene (Fig. 6B).

## Data availability

All data described in the manuscript are available in the GitHub repository: https://github.com/MaxTelford/Gainsandlosses.

## Code availability

All code described in the manuscript are available in the GitHub repository: https://github.com/MaxTelford/Gainsandlosses.

## Acknowledgements

We are grateful to Richard Copley for pointing out the possibility that errors underlie the phylogenetic signal in presence/absence data matrices and to members of the Telford lab for helpful comments. This work was funded by the BBSRC grant BB/R016240/1 (PK/MJT); by the European Union’s Horizon 2020 research and innovation programme under the Marie Skłodowska-Curie grant agreement No 764840 IGNITE (PN/MJT), and by the European Research Council grant (ERC-2012-AdG 322790) (PHS/MJT).

## Author information

### Affiliations

PN, PK, PHS, MJT: Centre for Life’s Origins and Evolution, Department of Genetics, Evolution and Ecology, University College London, London WC1E 6BT, UK. PHS current address: Institut für Zoologie, Sektion Entwicklungsbiologie, worm~lab, Universität zu Köln, Biozentrum Köln, Zülpicher Str. 47b, 50674 Köln, Deutschland

### Author contributions

Initial concept: MJT, PK, PN, PHS. Analyses: PN. Initial draft of manuscript: MJT, PN. Figures: PN. Final draft of MS: MJT, PN, PK, PHS.

---

**Supplementary Figure 1.**
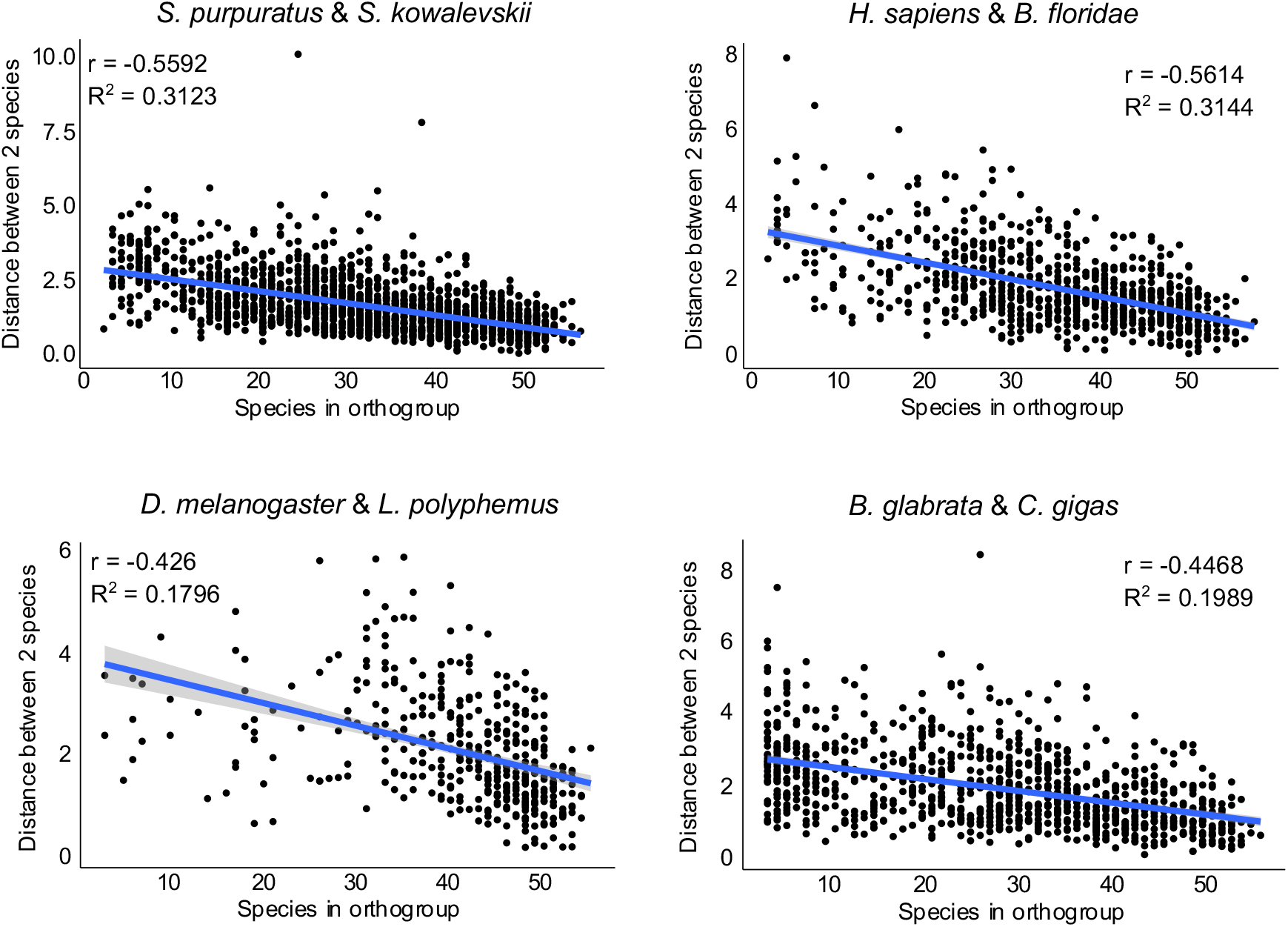
Faster-evolving genes are found in orthogroups with fewer taxa for empirical data. Each dot represents an orthogroup in which both species of the pair indicated are present. The data are from real genomes. The x axis shows the total number of different species present in the orthogroup and the y axis shows the patristic distance between the two species of the pair in the orthogroup tree (a measure of the rate of the gene). There is a small negative correlation trend between the two variables as shown by the Pearson’s correlation coefficient (*r*) and the adjusted R-squared of linear regression analysis (*R*^2^).

